# Onset of α5GABA-A Receptor Dependent Hippocampal Trisynaptic Circuit Dysfunction Is Associated Increased Age and Blood Pressure

**DOI:** 10.1101/2024.06.20.599963

**Authors:** Marcia H. Ratner, Richard Wainford, David H. Farb

## Abstract

Hypertension onset with aging is of widespread clinical significance, predominantly in males, yet the neural circuitry underpinnings for hypertension associated memory dysfunction remains unknown. Sprague Dawley (SD) male but not female rats develop age dependent increases in mean arterial blood pressure (MAP) by 16 months of age. We sought to interrogate the functional integrity of the hippocampal trisynaptic circuit (HTC), which is known to participate in memory, to determine whether age-associated increases in MAP contributes to circuitry dysfunction that may lead to mild-cognitive impairment (MCR). Ripples, and specifically sharp-wave ripple oscillations, play a role in memory replay and consolidation during awake immobility among other behaviors. These synchronous high frequency local field potentials (LFPs) in the ripple band (140 to 200 Hz) serve as an HTC level surrogate marker for circuitry function in rodents, non-human primates, and humans. Thus, we asked whether age-associated increased MAP might alter ripple dynamics. Recognizing that each patient responds in a unique way to hypertension we used a within subject design wherein each animal served as its own control in the investigative model. We surgically implanted high density silicon probe electrodes in HTC CA1 of young and aged SD males to determine whether a nootropic drug, α5IA, a negative allosteric modulator of α5 subunit containing type-A GABA receptors, could detect aberrant modulation of ripples within each subject. Here we report that acute oral administration of α5IA selectively modulated ripple amplitude, but not its duration or frequency during epochs of awake immobility. The response of peak ripple amplitude to α5IA is substantially diminished when chronic MAP exceeds 160 mmHg, corresponding to significant hypertension. The results are consistent with a model in which age-associated increases in MAP is associated with dysfunctional α5 GABA-A receptor modulation of ripple amplitude, but not duration or frequency, as a potential precision biomarker for memory dysfunction.

**Summary:** Age-related neurogenic hypertension disrupts memory and ripple band neural circuitry function in the hippocampal trisynaptic circuit involved in memory consolidation.

## INTRODUCTION

Untreated hypertension can lead to memory decline due to damaged brain vasculature, which can reduce blood flow and atrophy of nervous tissue, ultimately leading to vascular dementia (Petrea et al., 2020; Tayler et al., 2024). Reducing mean arterial blood pressure (MAP) with antihypertensives and lifestyle changes can impede degenerative changes (Affleck et al., 2023; Dao et al., 2024), but memory dysfunction can continue even with long-term pharmacotherapy. Moreover, outcomes are sensitive to the medication used, dosage, age at initiation of medication, and many aged individuals with hypertension will go on to develop dementia even while under antihypertensive control of MAP (den Brok et al., 2021). These clinical findings in human point to the significance of developing precision medicine approaches due to between subject variables. Within subject scientific research will hopefully provide a basis to identify a causality between biomarkers for hypertension, pharmacotherapy and biological outcomes.

The effects of chronic hypertension on neural network activity underlying memory have not, to the best of our knowledge, been previously investigated. The male Sprague Dawley (SD) rat develops increased MAP and impairments in memory function with aging, making it an excellent animal model for probing the role of hypertension in age-related amnestic mild-cognitive impairment (aMCI) (Dunnett et al., 1988; Johnson et al., 2020; Gannon et al., 2024). Synchronous high frequency activity in the ripple band (140 to 200 Hz) or sharp wave ripples (SPW-Rs) recorded from the CA1 hippocampal subregion is an established functional correlate of memory replay and consolidation during sleep and awake immobility in rodents, non-human primates, and humans (Buszaki et al., 1992 & 2015; Jadhav et al., 2012; Liu et al., 2022). SPW-R-associated IPSCs, but not EPSCs, are phase-locked with ripple cycles, indicating that phasic GABAergic interneurons play a role in the modulation of SPW-R oscillations (Gan et al., 2017). Based upon this phenomenon, we probed the hippocampal trisynaptic circuit (HTC) with a selective α5GABA-A receptor negative allosteric modulator (NAM) to ask whether the shaping of ripple oscillations in CA1 might be altered by MAP.

Using a within subject experimental design combined with oral administered drug, we previously demonstrated that α5IA, a nootropic, increases ripple band power and peak ripple amplitude in adult Fisher F344 and Long Evans male rats. However, the ripple bands of 9 to 16 mo TgF344-AD rats were unresponsive to α5IA, consistent with a role for α5GABAA receptor mediated inhibition in prodromal AD (Ratner et al., 2021). We hypothesized that if MAP and aging-induced memory dysfunction were part of AD and related dementias then high MAP might similarly disrupt HTC network function and becoming non-responsive to α5IA potentiation of the ripple band. We therefore investigated the relationship between age-related increases in MAP and HTC function using chronically implanted high-density silicon probes in HTC CA1 combined with *in vivo* electrophysiology. We observed that age-associated increased MAP is associated with substantial modulation of the amplitude of high frequency ripple band oscillations. These results strongly suggest that the neural network engaged in memory replay and consolidation is adversely affected by age-related MAP. This also indicates an opportunity, in future studies, to further understand the complex relationship between hypertension, MAP, aging, neural network pharmacology, and cognitive health.

## METHODS

### Subjects

Six male Sprague Dawley (SD) rats were used in these within subject *in vivo* electrophysiological experiments (**Table 1**). All rats were individually housed in a climate-controlled vivarium maintained on a regular 12hr/12hr light/dark cycle in the Laboratory Animal Science Center at the Boston University Chobanian & Avedisian School of Medicine School of Medicine. Rats had ad libitum access to water but were mildly food deprived to 85% of their free-feeding weight during training and testing. Rodent housing and research were both conducted in strict accordance with the NIH Guide for the Care and Use of Laboratory Animals. Boston University is accredited by the Association for Assessment and Accreditation of Laboratory Animal Care. The Boston University Institutional Animal Care and Use Committee approved all procedures described in this study.

**Table 1:** Baseline MAP, Age at *In Vivo* Electrophysiological Testing and Response to Probe Drug Challenge in male SD Rats.

| **Rat Number** | **MAP** | **Age at time of**<br>***In Vivo* Electrophysiological Testing**<br>**(Months/Days)** | **Response to α5IA Probe**<br>**Drug Challenge** |
| --- | --- | --- | --- |
| Young SD 1 | N/R | 5/0 | Responder |
| Young SD 2 | N/R | 3/17 | Responder |
| Aged SD 3 | 142 | 14/26 | Responder |
| Aged SD 4 | 152 | 15/15 | Responder |
| Aged SD 5 | 163 | 16/19 | ***Non-responder*** |
| Aged SD 6 | 183 | 16/29 | ***Non-responder*** |

N/R not recorded.

### Blood Pressure Measurement

Aged male SD rats are prone to developing increased MAP (the mean of systolic and diastolic blood pressure). Four aged male SD rats were tested blind for MAP prior to surgical placement of the silicon probe microelectrode arrays. MAP was measured with the non-invasive tail-cuff method using a WPI MRBP system (World Precision Instruments, Sarasota, FL, USA).

### Surgical Procedures

Rats were anesthetized with isoflurane (3.5% for induction; 1.5% - 2% thereafter) in 100% oxygen delivered via a calibrated vaporizer (Vaporizer Sales & Services Inc., Rockmart, Georgia, USA). The heads of the animals were shaved and prepped for surgery with betadine in triplicate. After placing the rats in a stereotaxic instrument (David Kopf Instruments, Tujunga, CA, USA), a midline sagittal incision was made across the top of the head to expose the cranium. A small craniotomy (∼2 mm in diameter) was created immediately above the right dorsal hippocampus with a center located at 3.6 mm AP and 2.5 mm ML from Bregma. The dura mater was excised with a dura knife, and the silicon microelectrode array lowered into position above the surface of the neocortex. Ten additional holes (< 0.5 mm) were drilled into the skull for the placement of an equal number of stainless-steel screws; two of these screws served as electrical grounds as well as providing additional anchoring points for securing the movable microelectrode array and copper mesh Faraday cage to the skull with dental acrylic (Patterson Dental Supply, St. Paul, Minnesota, USA) (Vandecasteele et al., 2012). The use of the Faraday cage in these experiments reduces electrical and radio frequency artifacts (Liu 2023). At the end of surgery, the silicon microelectrode array (NeuroNexus, Inc. Ann Arbor, Michigan, USA) was connected to an Intan RHD system (Intan, Inc) for positioning of the probe within the CA1 subregion based on real-time neural input. Electrode placement within the CA1 subregion were also adjusted after recovery from surgery and confirmed daily prior to training and test sessions by inspection of the local field potential (LFP) for evidence of theta activity during periods of ambulation and ripples during periods of immobility (Buzsáki 2022; 2015). These two neural activity metrics are well-established functional real-time indicators of electrode localization within the CA1 hippocampal subregion (Buzsáki 2022; 2015). Single shank silicon probes with linear electrode arrays that span across CA1 layers (str. oriens, pyramidal layer and str. radiatum) were also used in two of the rats included in these experiments to ensure that SPW-Rs are confidently measured and to facilitate rejection of artifacts due to chewing and movement (Liu 2022).

### Oral Drug Administration

The GABA-A α5-selective positive allosteric modulator α5IA (3-(5-methylisoxazol-3-yl)-6-[(1-methyl-1,2,3-triazol-4-yl) methyloxy]-1,2,4-triazolo[3,4-a] phthalazine) (Sigma Aldrich Inc, USA and Tecoland, Irvine, California, USA) was chosen based on its unique pharmacodynamic profile, oral bioavailability and our previous results demonstrating that it dose-dependently increases ripple band power and peak ripple amplitude in rats (Ratner et al., 2021). The vehicle (0.5% 400 centipoise methylcellulose) without or with drug were administered via oral gavage 30 min before each test session. To facilitate oral administration, rats were mildly sedated by briefly placing them in an induction chamber filled with 5% isoflurane. Drug was always administered after vehicle; counterbalancing the order of vehicle and drug administration was not possible due to the duration of the experimental protocol, the plasma half-life (0.9 hr) of the drug and, the within subject design for testing responsivity to a5IA.

### Escalating Cumulative Dose Paradigm for Analysis of Ripples

These experiments used our established probe drug challenge model which combines escalating doses of a5IA with serial recordings of local field potentials from the CA1 subregion while the animals are awake and immobile in a familiar environment (Ratner et al., 2021). This method was selected because different behaviors, such as running, sleeping and wakeful resting influence the occurrence of ripples in the CA1 hippocampal subregion. During the training phase, rats were allowed to explore a black plywood square (60 cm x 60 cm) enclosure with vertical yellow stripes on one of the four walls for at least 7 days to establish this as a “familiar environment”. Serial 10 min recordings were made while rats were immobile resting quietly in this familiar environment. Because electrode depth can influence power in the ripple band, the electrodes were not moved on the day of testing. For each series of experiments, we used single electrodes on each shank with optimal positioning in the region of interest to optimize comparisons within subject.

On probe drug testing day the subject is first placed in the same familiar environment and recording is initiated when exploratory ambulation ceases as judged by immobility while remaining awake and resting quietly. After each 10 min Familiar (F) recording session the subject is placed in small box on a turn table for 3 minutes. The environment is cleaned with 70% ethanol solution to remove odors. The rat is then returned to the same familiar (F) environment or placed in a Novel (N) environment to assess the behavioral and circuitry effect of novelty as called for by the protocol. The standard escalating dose protocol is: vehicle F1; dose-1 F2; dose-2 F3; dose-3 F4, where the dose listed in each figure represents the cumulative dose of drug at the time of each measurement. The same environment F is used in all 4 presentations, where F1 is the first exposure of the subject to the familiar environment and F4 is the 4^th^ exposure to the familiar environment.

### Data Acquisition and Post Acquisition Processing for Analysis

Three of the four aged SD rats were implanted with B32 type four shank silicon probes for effectively targeting a single subregion such as CA1 (NeuroNexus, Inc. Ann Arbor, Michigan, USA). Young SD 1 and aged SD 5 were both implanted with NeuroNexus A1X32 style single long shank probes which span multiple regions (e.g., CA1 and radiatum) allowing for signal comparison and ensuring the observations seen in these investigations are not artifacts of probe position. These probes also allowed us to determine electrode locations based on the distinct oscillatory patterns observed in different hippocampal subregions such as the hippocampal fissure based on theta power and the CA1 subregion based on the presence of ripples (Buszaki 2022). Our laboratory as well as others, have previously demonstrated that region specific functional signatures can be effectively used for real-time prediction of electrode localizations (Ratner et al., 2021; Ratner and Farb, 2022). Neural signal data was acquired using an Intan RHD system (Intan Technologies, Los Angeles, CA, USA) and processed using [NeuroExplorer](https://neuroexplorer.com/).

Electric fields monitored with extracellular electrodes provide useful information about state-dependent neuronal activity. The local field potential (LFP) recorded with indwelling microelectrodes represents the sum of the synchronized local changes in post-synaptic potentials measured in millivolts (mV). The greater the distance of the recording electrode from the source of these voltage changes, the less accurately the LFP reflects the activity interest (e.g., activity in the CA1 pyramidal cell layer) (Buszaki et al., 2012). The characteristics of the LFP signal include frequencies and amplitudes which reflect the changes in current flux associated with excitatory and inhibitory post synaptic potentials (Yochum et al., 2019). The LFP signal can be frequency filtered to allow for studying activity within specific frequency ranges. The amplitude of the LFP signal can be divided into its mean and peak components. A power spectral density (PSD) plot of the LFP signal can be generated by squaring the amplitude (mV) at each frequency (Hz). This plot reflects the relationship between the amplitude of the extracellular signal and its temporal frequency. Analysis of data from an LFP signal that has been frequency filtered for activity in the ripple band (140 to 200 Hz) provides information about the characteristics of the ripple events that give rise to both the raw LFP and the PSD plot of activity in this same frequency range. Thus, the mean and peak amplitude and frequencies of these events are reflected in the shape of the PSD plots. For this reason, we present both the raw data as a PSD plot and the computationally disambiguated peak ripple amplitude, mean ripple amplitude, ripple frequency, and ripple duration to improve the clarity of the neurophysiological findings.

Ripple event metrics were detected and characterized using the well-established method of the [Buzsaki Laboratory](https://github.com/buzsakilab/buzcode/blob/master/detectors/detectEvents/bz_FindRipples.m). Briefly, the LFP data was band pass frequency filtered for ripple band activity from 140 to 200Hz using a Butterworth filter. An event peak threshold, of 5.0 SDs of the band filtered LFP was used to identify potential ripple oscillations. Two additional thresholds defined as greater and less than 2.0 SDs of the filtered trace were used to define the beginning and end times for each ripple event. Ripples occurring within less than 30 ms of each other were merged into a single event. Only events with durations greater than 20 ms and less than 100 ms were included in the final analyses for significant differences in the averages of the mean and peak ripple amplitudes, peak frequencies, and durations or ripple events across the recording sessions. To ensure data reproducibility and improve rigor and transparency a commercially available program ([NeuroExplorer](https://www.neuroexplorer.com/downloads/NeuroExplorer5Manual.pdf)) was used to detect and process ripple events and calculate the ripple event metrics used in these studies. This commercially available platform was selected for this purpose with the goal of increasing the translational value of ripple data acquired by different laboratories (Ratner and Farb, 2022; Liu et al., 2022).

### Statistical Analysis

The processed ripple metric data was saved as a comma separated values (.csv) file and imported into GraphPad Prism for statistical analysis. Ripple metrics analyzed included: peak and mean ripple amplitudes (mV); peak ripple frequency (Hz); and ripple duration (sec). Appropriate non-parametric tests including Friedman’s ANOVA and the Kruskal-Wallis test were used for all within subject comparisons of ripple metrics. Student’s T test was used for between subject comparisons of average percent change in peak ripple amplitude in young SD versus aged hypertensive SD rats. Linear regression was used to assess for relationship between MAP and ripple metrics. Comparisons of ripple band power were performed on Z score normalized data sets using the Kolmogorov-Smirnov and Wilcoxon tests. Within and between subject comparisons of ripple band power using the 1.0 mg/kg dose was used for some within and between subject analyses because the occupancy percentage and variance at this dose are more favorable than those associated with a lower dose and higher doses in healthy rats without disease. This dose was also expected to have fewer off-target effects than the higher 2.0 mg/kg dose (Ratner et al., 2021).

## RESULTS

### MAP in Aged SD Male Rats

The MAPs of the aged male SDs were measured using a WPI MRBP tail-cuff system. The MAPs of the aged SDs included in this study ranged from 142 to 183 mmHg with a mean of 160 mmHg. **(Table 1)**

### MAP and Response to α5IA Probe Drug Challenge

*Within-subject analysis of the results of in vivo* electrophysiological recordings of high frequency neural activity in the ripple band also revealed *responders* and *non-responders* to α5IA probe drug challenge. Of the SD rats tested (n = 6), 33% were non-responders. Among the aged SD rats 50% were non-responders. This change in responder subtype was associated with a MAP greater than 160 mmHg which has previously been defined as severe hypertension in rats (Li et al., 2019).

### Power Spectral Density Plots Reveal Changes in Characteristic Neural Activity Patterns

Visual inspection of the Z-score normalized frequency filtered power spectral density (PSD) plots of the LFPs from young and aged rats reveals that age-associated increased MAP is associated with a conspicuous absence of the well-recognized characteristic hump in the ripple band (Sullivan et al., 2011; Buzsáki 2015; Ratner et al., 2021). **(Figs 1 A & B).** Analysis of the merged 1.0 mg/kg data from the young rats (n=2) and aged SDs (n = 4) reveals marked attenuation of the response to α5IA probe drug challenge in the aged SDs (**Fig 1C**).

**Figure 1:**
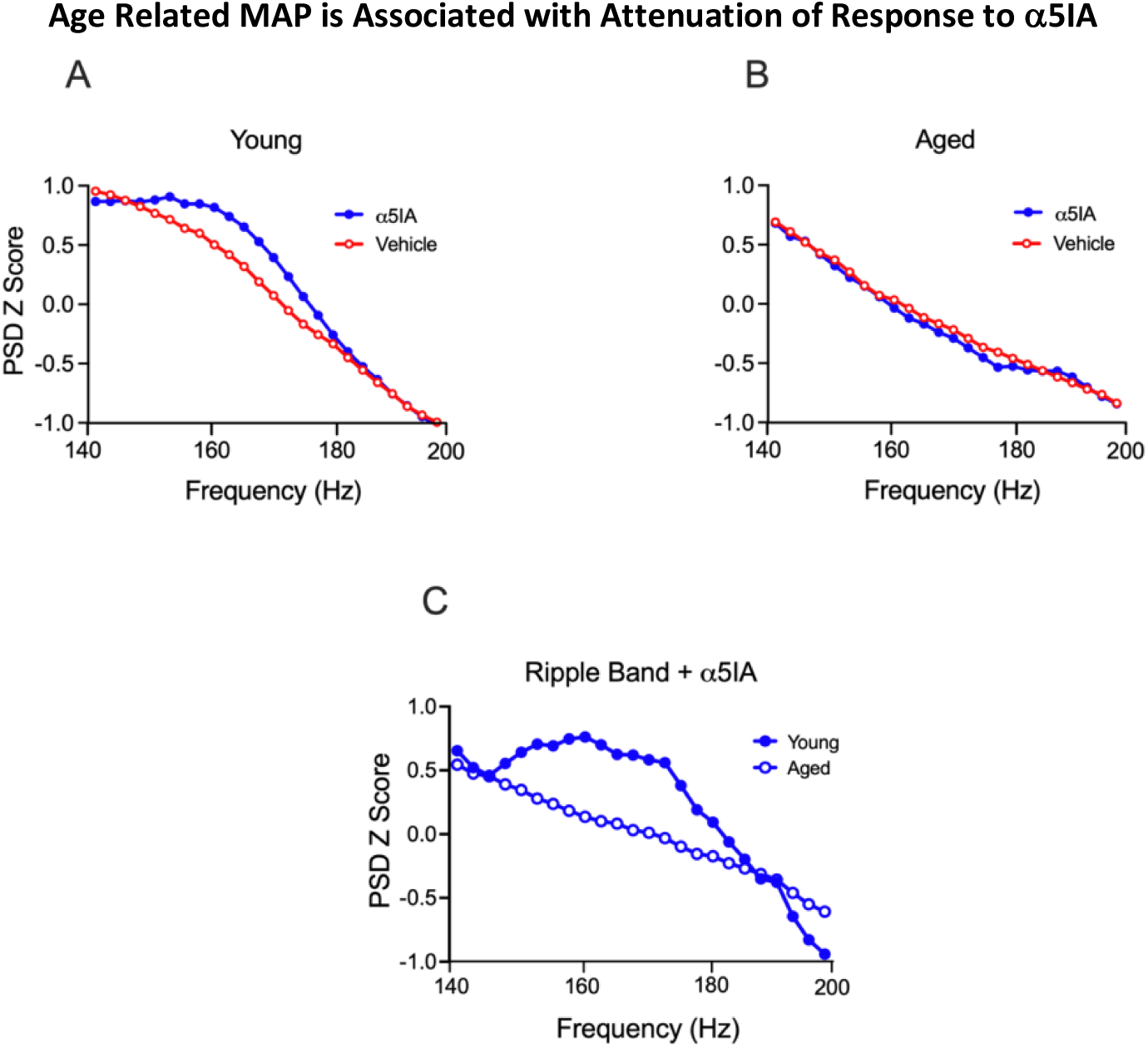
Age-associated High MAP Disrupts the Ripple Band. Power Spectral Density (PSD) Z score *normalized plots* on the ordinate axis with frequencies in Hz on the abscissa following administration of vehicle and 1.0 mg/kg of a5IA probe drug**. A)** PSD plot of a young SD rat showing significant increase (Wilcoxon test, p = 0.0006) in ripple band power in response to a5IA probe drug challenge with an overt hump in the ripple band. **B)** PSD plot from an aged high MAP SD (MAP 183 mmHg) reveals a conspicuous absence of the characteristic ripple band hump and small but significant (p = 0.001) attenuation in the response to a5IA probe drug challenge. C) Between subject analysis of Z score adjusted PSD plots of the merged 1.0 mg/kg data from the young rats (n=2) and aged SDs (n = 4) reveals a significant difference in the response to probe drug challenge in young and aged SDs (Kolmogorov-Smirnov, p = 0.002).

### Raw Power Spectral Density (PSD) Plots for all SD Rats

The raw power spectral density (PSD) plots frequency filtered for ripple band activity can be inspected for the presence or absence of the well-recognized characteristic hump from 140 to 200 Hz (Buzsáki 2015; Ratner et al., 2021; Ratner and Farb, 2022) following administration of 1.0 mg/kg a5IA, which produces increased ripple band power in adult F344 and Long Evans male rats (Ratner et al., 2021). This characteristic hump which is seen in young SDs and the aged SDs with a MAP under 150 mmHg is not seen in the older SDs which have progressively higher MAPs ranging from 152 to 183 mmHg. **(Fig 2).**

**Figure 2:**
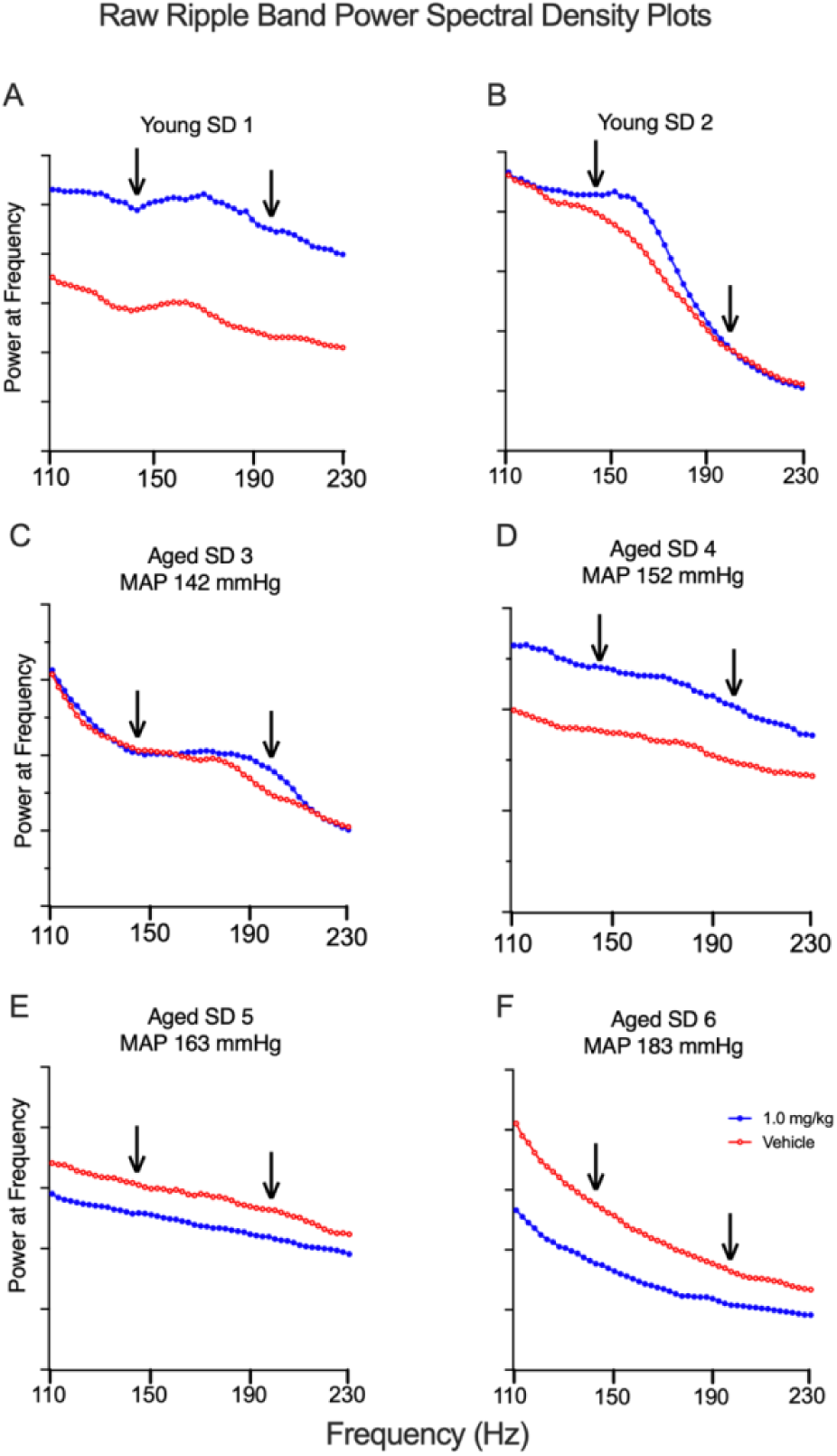
Raw Power Spectral Density (PSD) Plots for all Rats. PSD plots show relative frequency filtered local field potential power in mV^2^/Hz on the ordinate axis with frequencies in Hz on the abscissa following administration of vehicle and 1.0 mg/kg of α5IA probe drug**. A and B)** PSD plots of young male SD rats both show an overt hump in the ripple band indicated by arrows and an increase in relative power over vehicle baseline following administration of α5IA probe drug. **C)** PSD plot of aged SD 3 with MAP of 142 mmHg also shows an overt hump in the ripple band that increases following administration of the 1.0 mg/kg dose of α5IA but the frequency distribution is shifted to right. **D)** PSD plot of aged SD 3 which had the second lowest MAP also shows an overt increase in ripple band power in response to probe drug challenge but, the characteristic hump in the ripple band is overtly attenuated. **E-F)** PSD plots from the two aged SD rats with severe hypertension associated with MAPs greater than 160 mmHg reveals a conspicuous absence of the ripple band hump and a loss of the increase in ripple band power typically seen in response to α5IA probe drug challenge. Key shown in Panel F.

### Relationship Between Peak Ripple Amplitudes and MAPs

Statistical analysis for within subject effects of a5IA probe drug challenge on peak ripple amplitudes revealed a dose-dependent increase in the young SD rats as with healthy adult F344 and LE male rats (Ratner et al., 2021). **(Figs 3A & 4).** By contrast, age-associated increased MAP as defined by a MAP greater than 160 mmHg (Li et al., 2019), as seen in aged SD 3, was associated with loss of the dose-dependent probe drug challenge induced increases in peak ripple amplitude as in TgF344-AD rats **(Fig 3B).**

**Figure 3:**
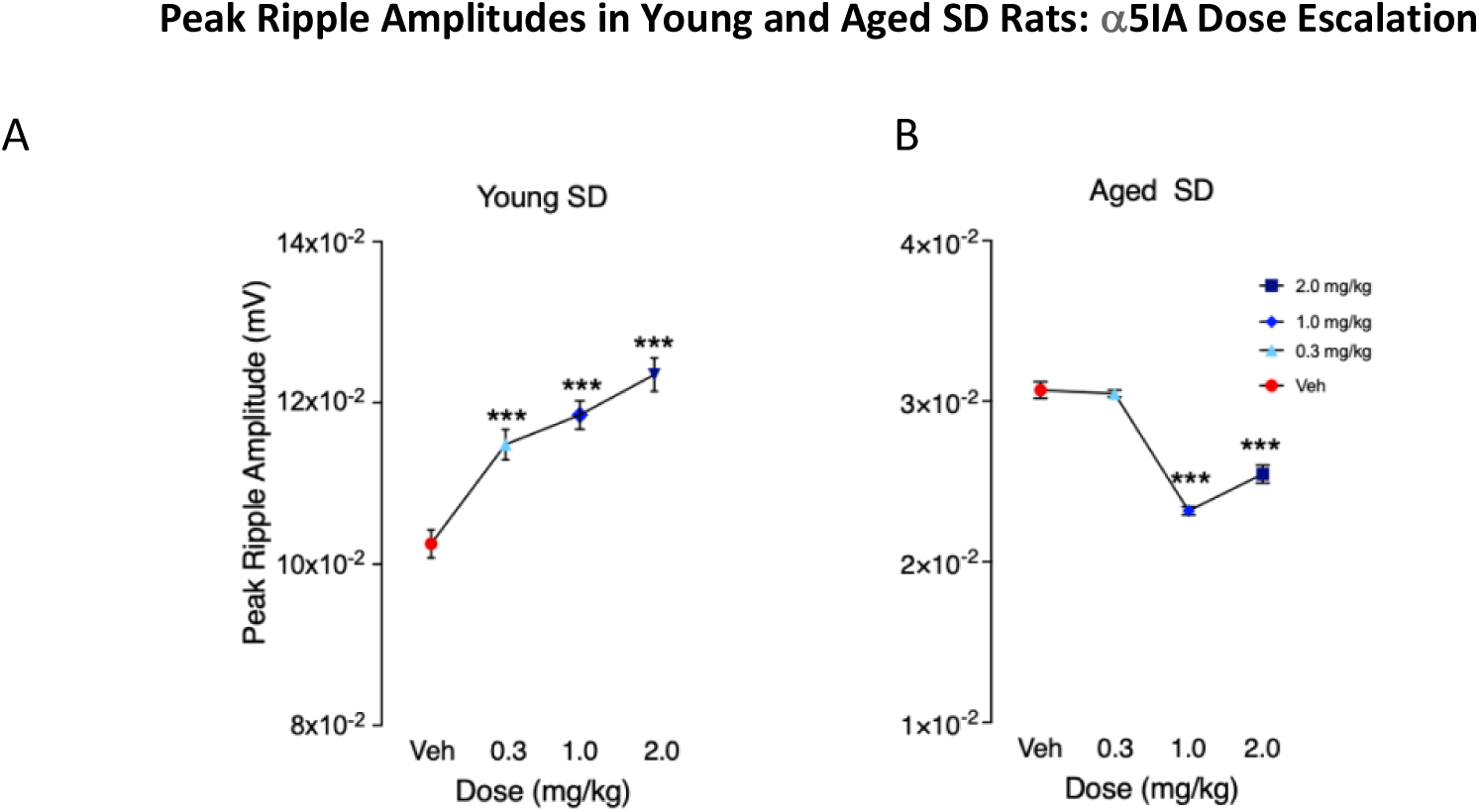
Representative Peak Ripple Amplitude Plots Showing Loss of Dose-Dependent Response to a5IA Probe Drug Challenge in aged SD. **A)** Plot from young male SD rat shows characteristic dose-dependent increases in peak ripple amplitudes following administration of all three doses of a5IA probe drug. All differences indicated as significant by asterisks are for drug dose compared with vehicle. The increase in peak ripple amplitude seen following the 2.0 mg/kg dose was also significantly greater than the 0.3 mg/kg dose (p = 0.007). **B)** Dose-response plot from an aged SD with severe hypertension (MAP > 160 mmHg) showing significant decreases in peak ripple amplitude following administration of the 1.0 and 2.0 mg/kg doses of a5IA. Results shown are from Kruskal-Wallis test with significant difference from vehicle indicated by *** at p < 0.001. Legend shown in Panel B.

### Peak Ripple Amplitudes Response to Probe Drug Challenge in SD Rats

Within subject analysis for the effects of α5IA probe drug challenge on peak ripple amplitudes revealed an increase in the two young SDs and the aged SDs with MAPs <160 mmHg which are similar to healthy adult male F344 and Long Evans rats (Ratner et al., 2021). **(Fig 4).** By contrast, aged SD Rats 3 & 4 which had MAPs greater than 160 mmHg show a loss of this probe drug induced increase in peak ripple amplitudes.

**Figure 4:**
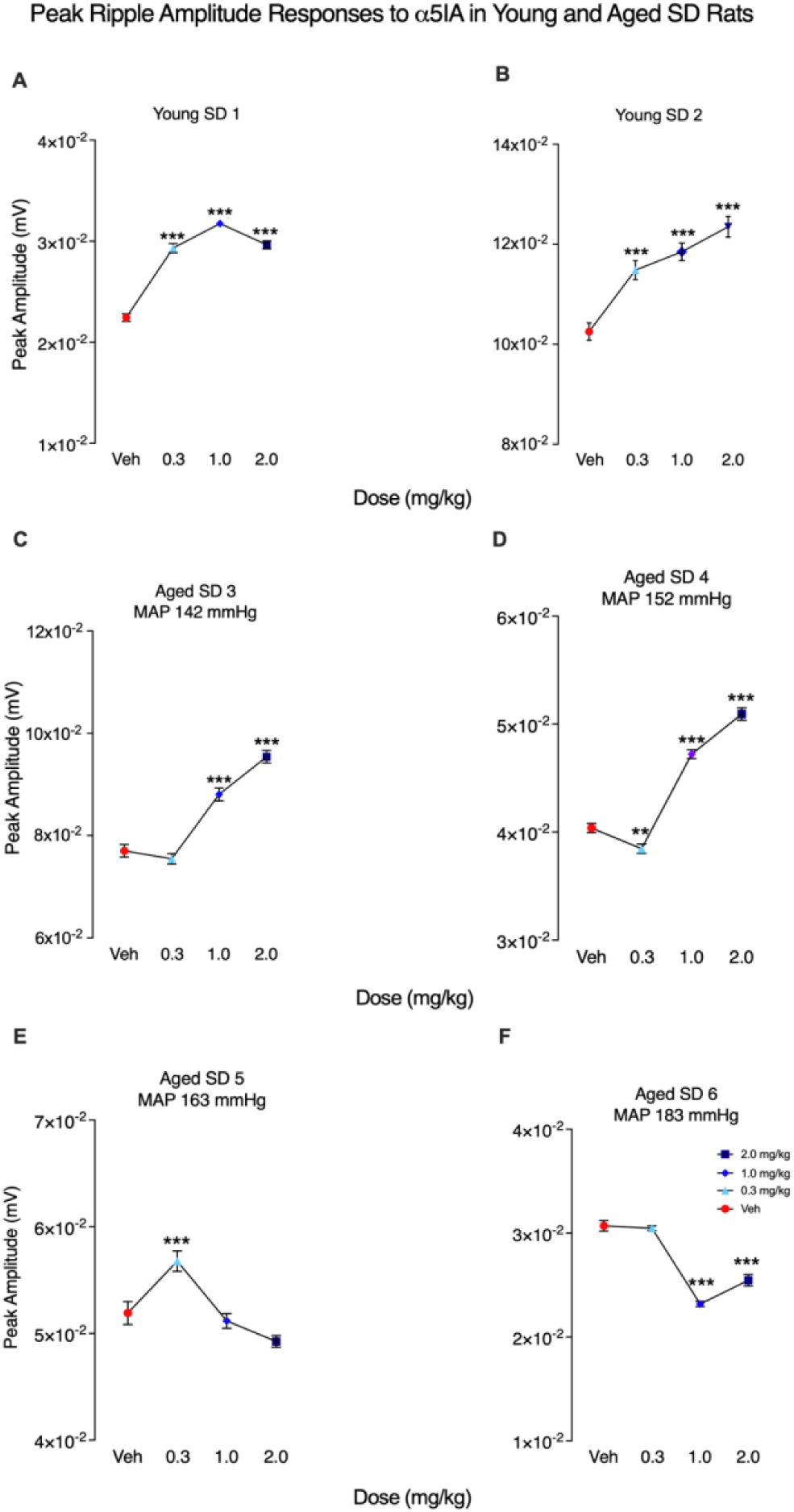
Average within-subject peak ripple amplitude responses to α5IA probe drug challenge. Plots show average dose-dependent within subject changes in peak ripple amplitude of all rats tested. **A-B)** Plots from young male SD rats showing characteristic increases in peak ripple amplitude following administration of all three doses of α5IA probe drug. **C-D)** Dose-response plots from aged SD 4 and 5 showing no change in average peak ripple amplitude at the lowest dose of α5IA followed by significant dose-dependent increases at the two higher doses. **E)** Dose-response plots from aged SD 6 showing a significant increase in average peak ripple amplitude following administration of the lowest dose of α5IA with no change at the two higher doses. **F)** Dose-response plots from aged SD 7 show no change in peak ripple amplitude at lowest dose test and decreases at the two higher doses. Results shown are from Kruskal-Wallis test with significance indicated by ** at p < 0.01 and **** at p < 0.0001. NS = not significant. Key shown in panel E.

Initial analyses were performed to look for average differences in peak ripple amplitudes both within groups and between groups. Analyses of the merged data from the young SD rats (n = 2) and the unstratified aged SD rats (n = 4) revealed that both groups responded to probe drug challenge with significant increases in peak ripple amplitude. **(Fig 5)** However, between group analysis of the merged raw data revealed that the aged SD rats had lower peak ripple amplitudes at all doses tested including vehicle baseline. **(Fig 5 & Table 2).**

**Figure 5:**
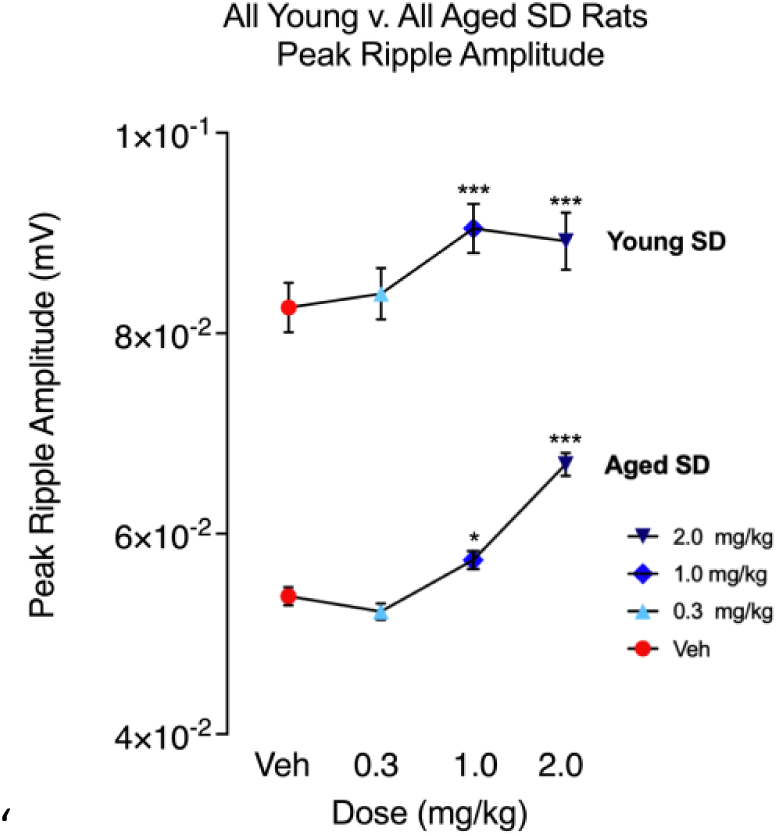
Average Peak Ripple Amplitude Responses to α5IA Probe Drug Challenge are Lower in all Young and Aged SD Rats. Plots show dose-dependent changes in averaged raw peak ripple amplitude values for young SD rats (n = 2) and unstratified aged SD rats (n = 4) following administration of escalating doses of α5IA. Significant results indicated by asterisks are from Kruskal-Wallis test with Dunn’s test for multiple comparisons for dose-dependent effects of α5IA versus vehicle with significance indicated by * at p < 0.05; *** at p < 0.001. In addition to the within group differences in the response to probe drug challenge shown above, significant between group difference in peak ripple amplitude were also observed (**Table 2**). Legend key for doses of α5IA shown in panel.

**Table 2:** Group Averaged Peak Ripple Amplitudes of Aged Male SD Rats are Significantly Lower than Young SD Rats at each Dose Tested in Probe Drug Challenge.

| α5IA Dose | Ripples Recorded Young vs. Aged | Young Mean ± SEM | Aged Mean ± SEM | Adjusted P Value* |
| --- | --- | --- | --- | --- |
| Vehicle | n<sub>Young</sub> = 277; n<sub>Aged</sub> = 551 | 0.08257 ± 0.002460 | 0.05307 ± 0.0009036 | <0.0001 |
| 0.3 mg/kg | n<sub>Young</sub> = 335; n<sub>Aged</sub> = 708 | 0.08394 ± 0.002552 | 0.05154 ± 0.0008059 | <0.0001 |
| 1.0 mg/kg | n<sub>Young</sub> = 360; n<sub>Aged</sub> = 767 | 0.09045 ± 0.002438 | 0.05667 ± 0.0008884 | <0.0001 |
| 2.0 mg/kg | n<sub>Young</sub> = 320; n<sub>Aged</sub> = 605 | 0.08919 ± 0.002852 | 0.06623 ± 0.001151 | 0.0010 |

\* Kruskal-Wallis Test with Dunn's Test for Multiple Comparisons; Young SD rats (n = 2); unstratified aged SD rats (n = 4); Total number of ripples (n = 3923)

Next, we plotted and compared the relative frequency distributions of peak ripple amplitudes in young and aged SD rats at vehicle baseline and following administration of escalating doses of α5IA to evaluate the changes in peak ripple amplitude response distribution that give rises to significant differences observed. This analysis revealed that the peak amplitude frequency distribution of the aged SD rats was shifted to the left as compared with the young SDs indicating that peak ripple amplitudes are lower in aged SDs (**Fig 6**). The relative frequency distribution of peak ripple amplitudes in the young SDs also shows an overt dose-dependent shift to the right in response to probe drug challenge with α5IA, a finding that is consistent with previously reports in F344 rats (Ratner et al., 2021).

**Figure 6:**
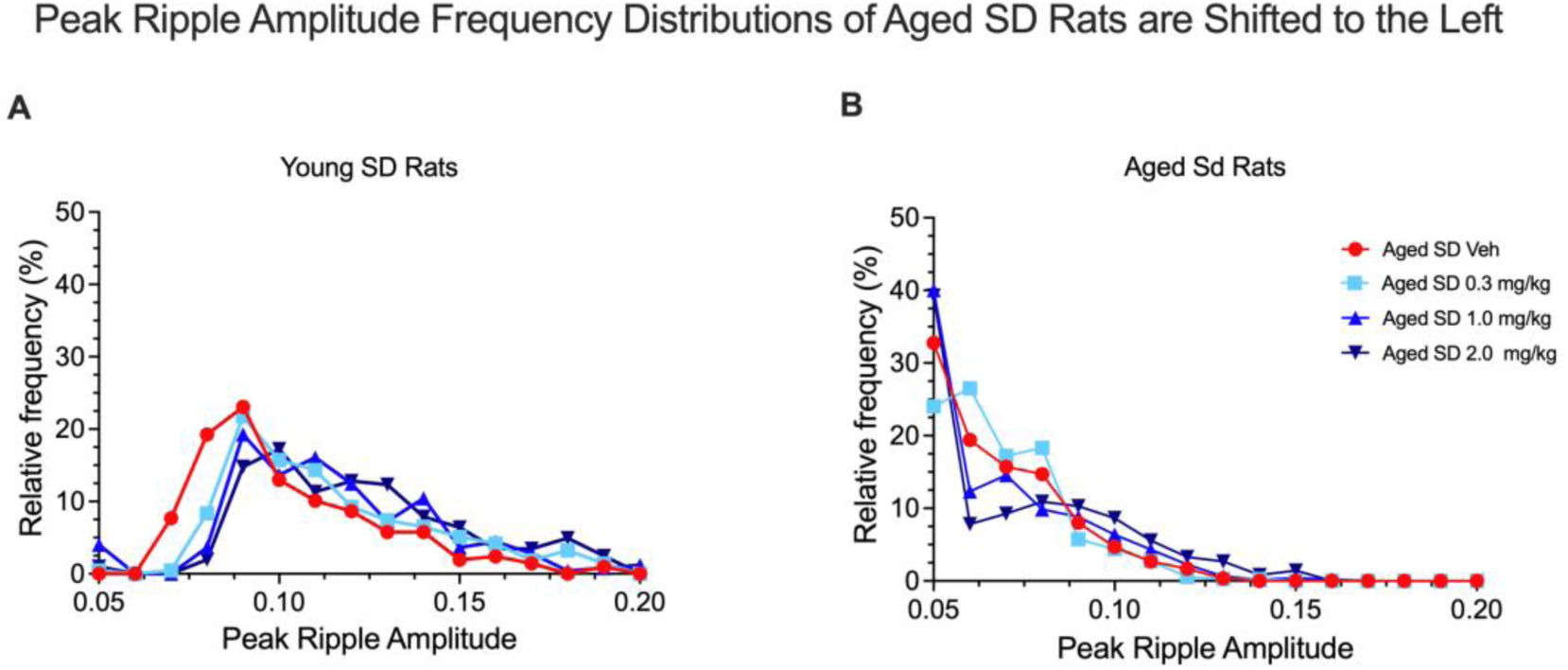
Peak Ripple Amplitude Frequency Distributions of Aged SD Rats are Shifted to Left. **A)** The average relative peak ripple amplitude frequency distributions for the two young SD rats following escalating doses of α5IA shows a drug-induced shift to the right with progressively higher doses of probe drug. **B)** The average relative frequency distribution for peak ripple amplitudes in the four aged SDs following administration of α5IA shows a distribution that favors lower amplitude events by comparison with the young SDs. Key shown in panel B.

Analysis of the merged data from the young SDs and aged SD rats stratified by MAPs revealed that the aged SDs with MAPs less than 160 mmHg responded to probe drug challenge with α5IA with an increase in peak ripple amplitude, while by contrast the aged SDs with MAPs greater than 160 mmHg did not respond to α5IA **(Fig 7).**

**Figure 7:**
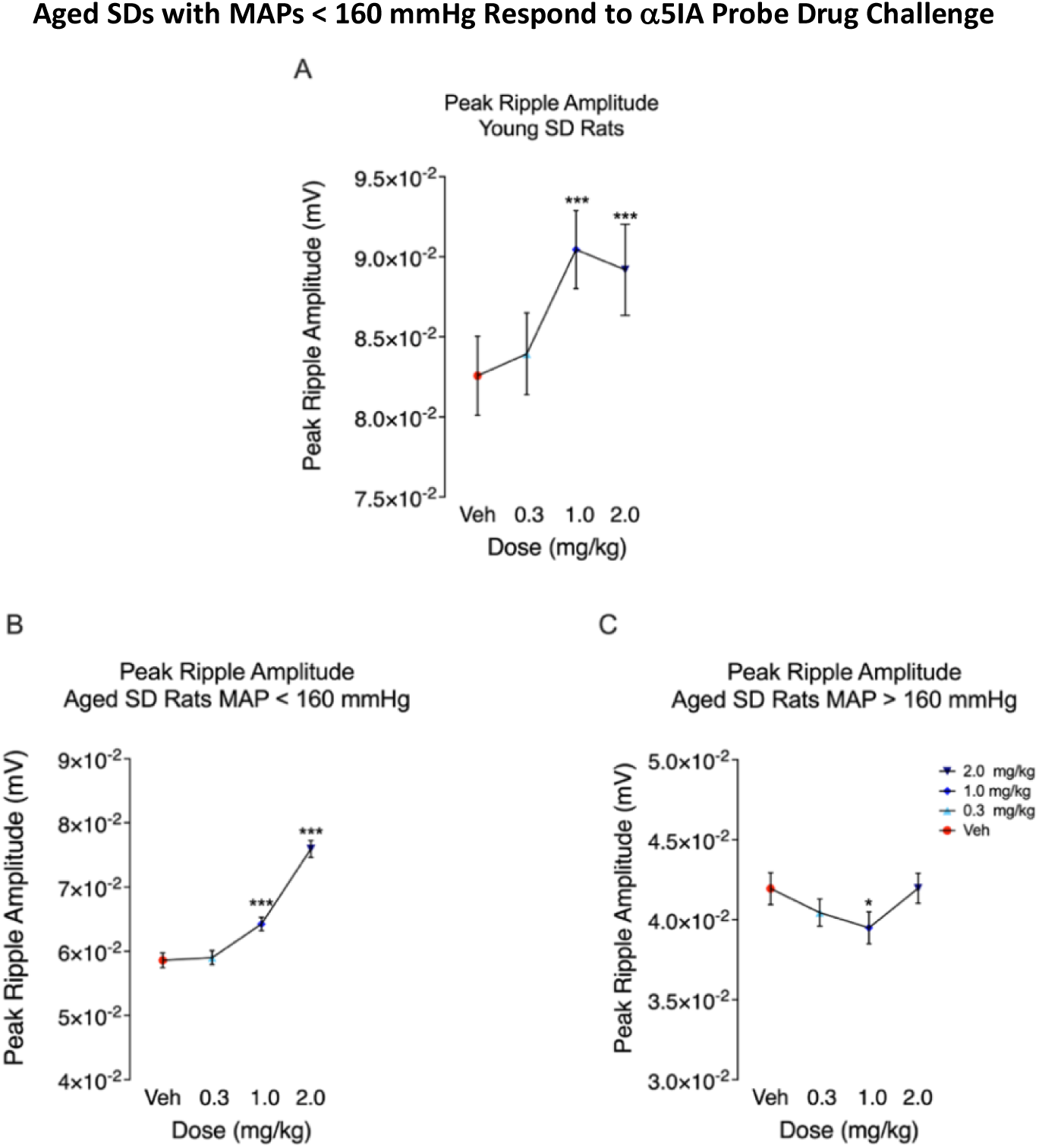
Aged SD Rats with MAPs greater than 160 mmHg do not Respond to Probe Drug Challenge with α5IA. **A)** Young SDs (n = 2) show significant dose-dependent increase in peak ripple amplitude following the 1.0 and 2.0 mg/kg doses of α5IA. **B)** Aged SDs with MAPs less than 160 mmHg (n = 2) show significant dose-dependent increase in peak ripple amplitude following the 1.0 and 2.0 mg/kg doses of α5IA. **C)** Aged SDs with MAPs greater than 160 mmHg (n = 2) only show a decrease in peak ripple amplitudes following the 1.0 mg/kg dose. Key shown in panel C.

To further quantify the relationship between age-associated increased MAP, the within subject peak ripple amplitudes were transformed to percent changes from a vehicle baseline set equal to zero (for ease of direct visual comparison). The data from the two young SDs was merged and served a control. These data were then plotted for visualization and analyzed for between subject effects. Significant differences in the average percent change in peak ripple amplitude was observed in the two aged SDs with age-associated increased MAP (MAP > 160 mmHg) **(Fig 8A)**. The corresponding descriptive statistics and results of a between Subject T tests comparing the average peak ripple amplitudes across all sessions from each of the aged SD rats with the averaged peak ripple amplitudes of the young SD rats are shown in **Table 3**. Results of linear regression revealed significant negative relationships between increasing MAP and percent change in peak ripple amplitude following administration of the 1.0 and 2.0 mg/kg doses of α5IA **(Fig 8B & Table 4).**

**Figure 8:**
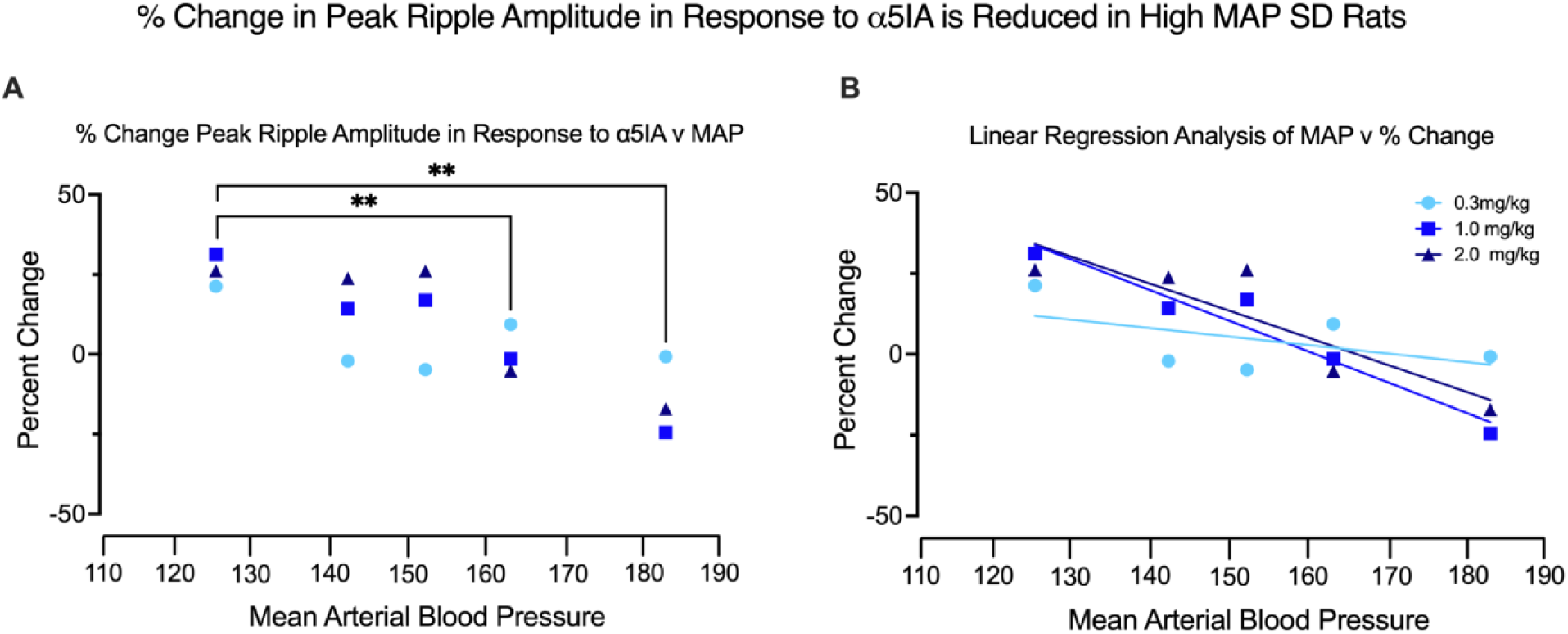
Lower Within-Subject Average Percent Change in Peak Ripple Amplitude Induced by Escalating Doses of α5IA is Associated with MAPs > 160 mm/Hg in Aged Male SD Rats. **A)** Graph shows percent change in peak ripple amplitude from 100% at vehicle baseline of zero induced by a5IA probe drug challenge at each dose tested in young and aged SD rats with progressively higher MAPs. Key shown in panel B. Significant differences from young SD rats are observed in the two aged SDs with severe hypertension characterized by a MAP greater than 160 mmHg (**Table 3**). **B)** Linear regression analyses of the relationships between MAPs and average percent change in peak ripple amplitude following escalating doses of α5IA in SD rats reveals significant relationships for the 1.0 and 2.0 mg/kg doses (**Table 4**).

**Table 3:** Aged SD Rats with Severe Hypertension have Significantly Lower Percent Changes in Average Peak Ripple Amplitude Following Probe Drug Challenge with α5IA.

| Rat | Mean Percent Change | Maximum | Minimum | Results of Student's T test |
| --- | --- | --- | --- | --- |
| Young Rats | 126.3 | 131.212573 | 121.300668 | N/A |
| Aged SD 1 | 112.8 | 126.126795 | 97.9615684 | p = 0.2322 |
| Aged SD 2 | 120.6 | 123.863931 | 95.2451709 | p = 0.1536 |
| Aged SD 3 | 100.9 | 109.381622 | 94.8757465 | **p = 0.0083** |
| Aged SD 4 | 85.9 | 99.32 | 75.496905 | **p = 0.006** |

**Table 4:** Linear Regression Reveals Significant Negative Relationships Between Peak Ripple Amplitude and MAPs in Aged SD Rats.

| α5IA Dose | 0.3 mg/kg | 1.0 mg/kg | 2.0 mg/kg |
| --- | --- | --- | --- |
| R<sup>2</sup> | 0.2798 | 0.9412 | 0.7880 |
| F | 1.165 | 48.04 | 11.15 |
| P value | 0.3594 | **0.0062** | **0.0444** |

## DISCUSSION

The results from our within-subject experiments comparing young to aged SD rats reveal previously unreported differences in high-frequency oscillations in the ripple band that are associated with age-related increases in MAP. We employed *in vivo* high density silicon probe electrophysiology with escalating doses of α5IA probe drug along with serial exposures to a familiar environment to try and modulate inhibitory interneuron shaping of SWRs in CA1 of the HTC by decreasing phasic and tonic synaptic inhibition. Local field potentials were recorded while animals were awake and immobile.

Aged male SD rats are known to develop increased MAP. By stratifying SD rats based on their responses to α5IA and MAP, differences in ripple band activity have been uncovered that were not apparent when the subjects were treated as a uniform group. These within subject MAP-dependent changes include: (1) the loss of a distinctive hump in the ripple band (which is characteristic of healthy young adult rats), (2) a shift of ripple amplitude distribution to lower amplitude oscillations, (3) a progressive decrease in the ability of α5IA to potentiate ripple amplitude, and (4) an apparent loss of the ability of α5IA to potentiate ripple amplitude with MAP > 160 mmHg.

These findings support the hypothesis that untreated age-associated increased MAP disrupts functional hippocampal circuitry that would likely underly performance deficits in aspects of spatial memory function. Some behavioral observations such as impaired performance on the delayed recognition trial of the Novel Object Recognition Test (NORT) are consistent with this reasoning (Dunnett et al., 1988; Johnson et al., 2020; Gannon et al., 2024). Additionally, age-associated increased MAP, either directly or as a comorbidity, disrupts high-frequency neural network oscillations in the ripple frequency band. The linkage between SPW-Rs, memory replay and consolidation were reviewed previously (Buzsáki, 2015).

The results of the α5IA modulation experiments support the hypothesis that tonic and/or phasic inhibitory control of CA1 ripple architecture results in increased peak ripple amplitude young adult SD animals but is disrupted in SD rats with severe age-associated increased MAP. The severity of hypertension correlates with greater functional deficits. Linear regression analyses indicate that the response to the 1.0 mg/kg dose of α5IA is the most sensitive indicator of ripple band dysfunction in this model. Interestingly, analyses of ripple event metrics from aged SD rats, which did not show the characteristic ripple band hump, nevertheless displayed a positive response to the probe drug challenge with significant increases in peak and mean ripple amplitudes. This suggests that rats with less severe age-associated increased MAP may initially retain some functionality, or the neural circuitry can compensate for the early adverse effects of hypertension. These in vivo electrophysiological findings in aged increased MAP SD rats are consistent with the hypothesis that untreated hypertension may lead to age-related neural network dysfunction implicated in aMCI and Alzheimer’s Disease (AD) (Dickstein et al., 2013; Pereira et al., 2014; Mahmmoud et al., 2015; Robitsek et al., 2015; Jones et al., 2019; Cowen et al., 2020).

CA3 population bursts, which give rise to ripples, are shaped by activity patterns in the dentate gyrus and entorhinal cortex, increasing the likelihood of ripples when dentate gamma power (50– 100 Hz) is optimal (Sullivan et al., 2011). These observations, combined with our findings of functional changes in the ripple amplitude distributions shifting to lower amplitude with age-associated increased MAP, suggests that upstream dysfunction in the entorhinal cortex, dentate gyrus, CA3 subregions, or some combination of them will almost certainly play a role in the qualitative differences observed in the PSD plots and the quantitative modulatory responses to α5IA. It is noteworthy that studies of 6-month-old spontaneously hypertensive Wistar rats have shown a reduction in grey matter volume in both the dentate gyrus and CA1 subregions, alongside a loss of neurons in the CA1 subregion (Sabbatini et al., 2000, 2002).

This study has several limitations. We did not record neural data from other hippocampal subregions, such as the dentate gyrus (DG), so we cannot determine whether changes in DG gamma power contributes to the observed changes in high frequency. Second, due to the risk of dislodging the microelectrode arrays, we were unable to measure MAP immediately prior to recording, and therefore, MAPs were measured before rats were implanted. Third, the behavioral aspects of this study are limited to the interactions between probe drug administration and periods of spontaneous immobility when theta power is low, and ripple events are more likely to occur. Therefore, the influence of α5IA probe drug challenge on neural activity patterns in the CA1 subregion associated with ambulation, such as theta-gamma coupling, could not be effectively investigated with this model (van den Berg et al., 2023).

## CONCLUSIONS

The results reveal qualitative and quantitative differences in ripple band activity due to age-associated increased MAP which cannot be explained by electrode depth. These changes include: 1) a loss of the characteristic hump seen on power spectral density plots of the ripple band; 2) decreased peak ripple amplitudes and a shift to the left in the frequency distribution of peak ripple amplitudes; 3) a loss of the response to probe drug challenge with a5IA in rats with severe age-associated increased MAP (MAP > 160 mgHg); 4) a 1.0 mg/kg dose of a5IA is the most sensitive indicator of ripple band dysfunction due to age-associated increased MAP. In conclusion, the results of these investigations indicate that high frequency neural activity in the ripple band which is implicated in synaptic plasticity and memory consolidation is sensitive to the adverse effects of age-associated increased MAP. These findings also suggest that control of MAP during aging might prevent the onset of changes in ripple band activity including a loss of the response to a5IA which are like those seen in the TgF344-SD rat model of Alzheimer’s disease (Ratner et al., 2021).

## ACKNOWLEDGEMENTS

The authors thank Alexis Muschal for assistance with the blood pressure measurements in the aged hypertensive rats included in this study.

## Notes

### Competing Interest Statement

The authors have declared no competing interest.

## REFERENCES

Affleck AJ, Sachdev PS, Halliday GM. Past antihypertensive medication use is associated with lower levels of small vessel disease and lower Aβ plaque stage in the brains of older individuals. Neuropathol Appl Neurobiol. 2023 Aug;49(4):e12922. doi: 10.1111/nan.12922. PMID: 37431095.

Buzsáki G. Theta oscillations in the hippocampus. Neuron. 2002 Jan 31;33(3):325–40. doi: 10.1016/s0896-6273(02)00586-x. PMID: 11832222.

Buzsáki G, Anastassiou CA, Koch C. The origin of extracellular fields and currents--EEG, ECoG, LFP and spikes. Nat Rev Neurosci. 2012 May 18;13(6):407–20. doi: 10.1038/nrn3241. PMID: 22595786; PMCID: PMC4907333.

Buzsáki G. Hippocampal sharp wave-ripple: A cognitive biomarker for episodic memory and planning. Hippocampus. 2015 Oct;25(10):1073–188. doi: 10.1002/hipo.22488. PMID: 26135716; PMCID: PMC4648295..

Cowen SL, Gray DT, Wiegand JL, Schimanski LA, Barnes CA. Age-associated changes in waking hippocampal sharp-wave ripples. Hippocampus. 2020 Jan;30(1):28–38. doi: 10.1002/hipo.23005. Epub 2018 Nov 11. PMID: 29981255; PMCID: PMC6322975.

Dao E, Barha CK, Zou J, Wei N, Liu-Ambrose T. Prevention of Vascular Contributions to Cognitive Impairment and Dementia: The Role of Physical Activity and Exercise. Stroke. 2024 Feb 27. doi: 10.1161/STROKEAHA.123.044173. Epub ahead of print. PMID: 38410973.

den Brok MGHE, van Dalen JW, Abdulrahman H, Larson EB, van Middelaar T, van Gool WA, van Charante EPM, Richard E. Antihypertensive Medication Classes and the Risk of Dementia: A Systematic Review and Network Meta-Analysis. J Am Med Dir Assoc. 2021 Jul;22(7):1386–1395.e15. doi: 10.1016/j.jamda.2020.12.019. Epub 2021 Jan 16. PMID: 33460618.

Dickstein DL, Weaver CM, Luebke JI, Hof PR. Dendritic spine changes associated with normal aging. Neuroscience. 2013 Oct 22;251:21–32. doi: 10.1016/j.neuroscience.2012.09.077. Epub 2012 Oct 13. PMID: 23069756; PMCID: PMC3654095.

Dunnett SB, Evenden JL, Iversen SD. Delay-dependent short-term memory deficits in aged rats. Psychopharmacology (Berl). 1988;96(2):174–80. doi: 10.1007/BF00177557. PMID: 3148143.

Gannon O, Tremble SM, McGinn C, Guth R, Scoppettone N, Hunt RD, Prakash K, Johnson AC. Angiotensin II-mediated hippocampal hypoperfusion and vascular dysfunction contribute to vascular cognitive impairment in aged hypertensive rats. Alzheimers Dement. 2024 Feb;20(2):890–903. doi: 10.1002/alz.13491. Epub 2023 Oct 10. PMID: 37817376.

Johnson AC, Miller JE, Cipolla MJ. Memory impairment in spontaneously hypertensive rats is associated with hippocampal hypoperfusion and hippocampal vascular dysfunction. J Cereb Blood Flow Metab. 2020 Apr;40(4):845–859. doi: 10.1177/0271678X19848510. Epub 2019 May 14. PMID: 31088235; PMCID: PMC7168795.

Jones EA, Gillespie AK, Yoon SY, Frank LM, Huang Y. Early Hippocampal Sharp-Wave Ripple Deficits Predict Later Learning and Memory Impairments in an Alzheimer’s Disease Mouse Model. Cell Rep. 2019 Nov 19;29(8):2123–2133.e4. doi: 10.1016/j.celrep.2019.10.056. PMID: 31747587; PMCID: PMC7437815.

Li J, Kemp BA, Howell NL, Massey J, Mińczuk K, Huang Q, Chordia MD, Roy RJ, Patrie JT, Davogustto GE, Kramer CM, Epstein FH, Carey RM, Taegtmeyer H, Keller SR, Kundu BK. Metabolic Changes in Spontaneously Hypertensive Rat Hearts Precede Cardiac Dysfunction and Left Ventricular Hypertrophy. J Am Heart Assoc. 2019 Feb 19;8(4):e010926. doi: 10.1161/JAHA.118.010926. PMID: 30764689; PMCID: PMC6405673.

Liu AA, Henin S, Abbaspoor S, Bragin A, Buffalo EA, Farrell JS, Foster DJ, Frank LM, Gedankien T, Gotman J, Guidera JA, Hoffman KL, Jacobs J, Kahana MJ, Li L, Liao Z, Lin JJ, Losonczy A, Malach R, van der Meer MA, McClain K, McNaughton BL, Norman Y, Navas-Olive A, de la Prida LM, Rueckemann JW, Sakon JJ, Skelin I, Soltesz I, Staresina BP, Weiss SA, Wilson MA, Zaghloul KA, Zugaro M, Buzsáki G. A consensus statement on detection of hippocampal sharp wave ripples and differentiation from other fast oscillations. Nat Commun. 2022 Oct 12;13(1):6000. doi: 10.1038/s41467-022-33536-x. PMID: 36224194; PMCID: PMC9556539.

Mahmmoud RR, Sase S, Aher YD, Sase A, Gröger M, Mokhtar M, Höger H, Lubec G. Spatial and Working Memory Is Linked to Spine Density and Mushroom Spines. PLoS One. 2015 Oct 15;10(10):e0139739. doi: 10.1371/journal.pone.0139739. PMID: 26469788; PMCID: PMC4607435.

Pereira AC, Lambert HK, Grossman YS, Dumitriu D, Waldman R, Jannetty SK, Calakos K, Janssen WG, McEwen BS, Morrison JH. Glutamatergic regulation prevents hippocampal-dependent age-related cognitive decline through dendritic spine clustering. Proc Natl Acad Sci U S A. 2014 Dec 30;111(52):18733–8. doi: 10.1073/pnas.1421285111. Epub 2014 Dec 15. PMID: 25512503; PMCID: PMC4284552.

Petrea RE, O’Donnell A, Beiser AS, Habes M, Aparicio H, DeCarli C, Seshadri S, Romero JR. Mid to Late Life Hypertension Trends and Cerebral Small Vessel Disease in the Framingham Heart Study. Hypertension. 2020 Sep;76(3):707–714. doi: 10.1161/HYPERTENSIONAHA.120.15073. Epub 2020 Jul 31. PMID: 32755403; PMCID: PMC7577565.

Ratner MH, Downing SS, Guo O, Odamah KE, Stewart TM, Kumaresan V, Robitsek RJ, Xia W, Farb DH. Prodromal dysfunction of α5GABA-A receptor modulated hippocampal ripples occurs prior to neurodegeneration in the TgF344-AD rat model of Alzheimer’s disease. Heliyon. 2021 Sep 1;7(9):e07895. doi: 10.1016/j.heliyon.2021.e07895. PMID: 34568591; PMCID: PMC8449175.

Ratner MH, Farb DH. Probing the Neural Circuitry Targets of Neurotoxicants In Vivo Through High Density Silicon Probe Brain Implants. Front Toxicol. 2022 Apr 25;4:836427. doi: 10.3389/ftox.2022.836427. PMID: 35548683; PMCID: PMC9081674.

Robitsek J, Ratner MH, Stewart T, Eichenbaum H, Farb DH. Combined administration of levetiracetam and valproic acid attenuates age-related hyperactivity of CA3 place cells, reduces place field area, and increases spatial information content in aged rat hippocampus. Hippocampus. 2015 Dec;25(12):1541–55. doi: 10.1002/hipo.22474. Epub 2015 Jul 14. PMID: 25941121; PMCID: PMC4633399.

Sabbatini M, Strocchi P, Vitaioli L, Amenta F. The hippocampus in spontaneously hypertensive rats: a quantitative microanatomical study. Neuroscience. 2000;100(2):251–8. doi: 10.1016/s0306-4522(00)00297-9. PMID: 11008165.

Sabbatini M, Catalani A, Consoli C, Marletta N, Tomassoni D, Avola R. The hippocampus in spontaneously hypertensive rats: an animal model of vascular dementia? Mech Ageing Dev. 2002 Mar 15;123(5):547–59. doi: 10.1016/s0047-6374(01)00362-1. PMID: 11796140.

Sullivan D, Csicsvari J, Mizuseki K, Montgomery S, Diba K, Buzsáki G. Relationships between hippocampal sharp waves, ripples, and fast gamma oscillation: influence of dentate and entorhinal cortical activity. J Neurosci. 2011 Jun 8;31(23):8605–16. doi: 10.1523/JNEUROSCI.0294-11.2011. PMID: 21653864; PMCID: PMC3134187.

Tayler HM, Skrobot OA, Baron DH, Kehoe PG, Miners JS. Dysregulation of the renin-angiotensin system in vascular dementia. Brain Pathol. 2024 Mar 7:e13251. doi: 10.1111/bpa.13251. Epub ahead of print. PMID: 38454306.

Vandecasteele M, M S, Royer S, Belluscio M, Berényi A, Diba K, Fujisawa S, Grosmark A, Mao D, Mizuseki K, Patel J, Stark E, Sullivan D, Watson B, Buzsáki G. Large-scale recording of neurons by movable silicon probes in behaving rodents. J Vis Exp. 2012 Mar 4;(61):e3568. doi: 10.3791/3568. PMID: 22415550; PMCID: PMC3399468.

van den Berg M, Toen D, Verhoye M, Keliris GA. Alterations in theta-gamma coupling and sharp wave-ripple, signs of prodromal hippocampal network impairment in the TgF344-AD rat model. Front Aging Neurosci. 2023 Mar 22;15:1081058. doi: 10.3389/fnagi.2023.1081058. PMID: 37032829; PMCID: PMC10075364.

Yochum M, Modolo J, Mogul DJ, Benquet P, Wendling F. Reconstruction of post-synaptic potentials by reverse modeling of local field potentials. J Neural Eng. 2019 Apr;16(2):026023. doi: 10.1088/1741-2552/aafbfb. Epub 2019 Jan 4. PMID: 30609420.

